# *Elongation of Very Long Chain Fatty Acids Like- 3* (*Elovl3*) is activated by ZHX2 and is a regulator of cell cycle progression

**DOI:** 10.1101/2022.09.02.506374

**Authors:** Kate Townsend Creasy, Hui Ren, Jieyun Jiang, Martha L. Peterson, Brett T. Spear

**Author notes:** Corresponding Author: Department of Microbiology, Immunology & Molecular Genetics, University of Kentucky Medical Center, Lexington, KY 40536-0096. Department of Biobehavioral Health Sciences, School of Nursing, University of Pennsylvania, Philadelphia, PA. Epocrates at athenahealth, Austin, TX.

## Abstract

Zinc fingers and homeoboxes 2 (ZHX2) functions as a tumor suppressor in several models of hepatocellular carcinoma (HCC), presumably through its control of target genes. Previous microarray data suggested that Elongation of Very Long Chain Fatty Acids 3 (Elovl3), a member of the Elovl family which synthesize very long chain fatty acids (VLCFAs), is a putative ZHX2 target gene. VLCFAs are core component of ceramides and other bioactive sphingolipids, which are often dysregulated in diseases, including HCC. Since several previously identified ZHX2 targets become dysregulated in HCC, we investigated the relationship between ZHX2 and *Elovl3* in liver damage and HCC. Here, using mouse and cell models, we demonstrate that *Zhx2* positively regulates *Elovl3* expression in the liver and that male-biased hepatic *Elovl3* expression is established between 4-8 weeks of age in mice. *Elovl3* is dramatically repressed in mouse models of liver regeneration and HCC and the reduced *Elovl3* levels in the regenerating liver are associated with changes in hepatic very long chain fatty acids. Human hepatoma cell lines with forced *Elovl3* expression have lower rates of cell growth; analysis of synchronized cells indicate that this reduced proliferation correlates with cells stalling in S-phase. Taken together, these data indicate that *Elovl3* expression helps regulate cellular proliferation, possibly through control of VLCFAs, and its repression may be a contributing factor to HCC and explain, in part, the function of ZHX2 as a suppressor of HCC progression.

## Introduction

Zinc fingers and homeoboxes 2 (*Zhx2*) is emerging as a critical transcriptional regulator of hepatic gene expression during development and disease. BALB/cJ mice, which have a natural hypomorphic mutation in the *Zhx2* gene,^1,2^ and engineered *Zhx2* knockout mice have altered expression of numerous liver genes. In the absence of *Zhx2*, genes that are normally silenced in the postnatal liver, including *alpha-fetoprotein* (*AFP*), *H19, glypican 3* (*Gpc3*), and *lipoprotein lipase* (*Lpl*), continue to be expressed.^1,3,4^ These Zhx2 target genes are frequently dysregulated during liver regeneration and in hepatocellular carcinoma (HCC).^5^ ZHX2 also influences expression of sex-biased target genes, including male-biased *major urinary protein (Mup)* and female-biased *cytochrome P450* (*Cyp*) genes;^6,7^ these genes are also dysregulated in regenerating liver. Furthermore, ZHX2 regulates hepatic lipid metabolism genes with notable effects on plasma lipid levels and consequent cardiovascular disease risk.^8^

We and others have proposed that ZHX2 functions as a tumor suppressor, presumably through the dysregulation of ZHX2 target genes.^5,9^ Several studies in HCC cells lines and mice support this possibility. ZHX2 blocks cell proliferation through direct binding to the promoters of Cyclins A and E and inhibiting their expression.^10^ In addition, ZHX2 inhibits expression of Lpl and activates miR-24-3P, which reduces SREBP1c levels; these activities block both lipid uptake and de novo lipogenesis and subsequent tumor progression in cell lines.^11^ Since numerous hepatic genes are dysregulated in the absence of ZHX2, additional ZHX2 targets may also contribute to HCC progression.

Liver microarray data from BALB/cJ and ZHX2-positive congenic mice showed a positive correlation between ZHX2 and Elongation of very long chain fatty acids-3 (*Elovl3)* levels.^8^ ELOVL3 belongs to a family of enzymes (ELOVL 1-7) that synthesize very long chain fatty acids (VLCFAs), complex lipids comprising fatty acids with 20 or more carbons (≥C20).^12^ VLCFAs are commonly incorporated into ceramides or metabolized to lipid mediators that can regulate cellular growth, differentiation, proliferation, and other physiological functions.^13^ ELOVL3, which is expressed primarily in brown and white adipose tissue, skin sebaceous glands, and the liver, synthesizes C20-C24 saturated and monounsaturated fatty acids.^14-16^

Dysregulated lipid metabolism is a key feature of numerous liver pathologies, including hepatocellular carcinoma (HCC).^17,18^ We therefore investigated a potential link between ZHX2, ELOVL3, and HCC. Using several mouse models and tissue culture assays, we demonstrate that ZHX2 is a transcriptional activator of hepatic *Elovl3* expression. Furthermore, consistent with altered expression of other ZHX2 targets, we find that *Elovl3* mRNA levels decrease dramatically in mouse models of liver regeneration and HCC. Consistent with this, hepatic concentrations of ELOVL3-synthesized VLCFAs are concomitantly lower in regenerating livers. We also investigated whether the changes in *Elovl3* expression might impact cell proliferation. Our studies reveal that forced *Elovl3* expression slows the growth and expansion of human hepatoma cells. Furthermore, studies in synchronized cells indicate *Elovl3* expression stalls the cell cycle in S-phase. These data suggest ZHX2 tumor suppressor activities may be enacted, at least in part, by enhanced *Elovl3* expression that inhibits cellular proliferation.

## Materials and Methods

### Mouse Strains

All mice were housed in the University of Kentucky Division of Laboratory Animal Research facility and maintained in accordance with Institutional Animal Care and Use Committee (IACUC) approved protocols. All mice had *ad libitum* access to food and water and were maintained on a 14/10-hour light/dark cycle. BALB/cJ and BALB/cByJ mice were obtained from The Jackson Laboratory. Hepatocyte-specific *Zhx2* knockout (*Zhx2*^Δ*hep*^) and whole-body Zhx2 knockout (*Zhx2*^*KO*^) mice have been previously reported. ^6,7^ Briefly, C57BL/6 (BL/6) mice homozygous with floxed *Zhx2* alleles (*Zhx2*^*f/f*^) were bred to BL/6 mice expressing Albumin promoter-driven Cre recombinase (*Alb-Cre*, Jackson Labs #003574) or BL/6 mice expressing E2a promoter-driven Cre recombinase (E2a-Cre, Jackson Labs #003724) and then back-crossed to produce hepatocyte-specific *Zhx2* knockout mice (*Zhx2*^Δ*hep*^) or whole-body *Zhx2*^*KO*^ mice, respectively. Littermates without Cre (*Zhx2*^*f/f*^), which express *Zhx2* at the same levels as wild-type mice, were used as controls.

### Developmental timepoint studies

Female C3H/HeJ (C3H) mice were bred to male BL/6 mice to generate C3B6F1 offspring; these mice were used since they have the wild-type Zhx2 gene and were used for studies to compare Zhx2 and Elovl3 mRNA levels. Pregnant mice were monitored for vaginal plugs then euthanized after 17 days to collect livers for embryonic timepoints. Neonatal pups and mature mice were euthanized by age-appropriate methods at the indicated timepoints. Livers were excised, snap frozen, and stored at -80°C until further analysis.

### Liver Regeneration

Liver regeneration was induced by a single intraperitoneal injection of carbon tetrachloride (CCl_4_). Adult male C3H, BL/6 and B6C3F1 mice used since C3H and B6C3F1 mice have higher AFP activation after CCl4 treatment than BL/6 mice, and we wanted to test whether Elovl3 also exhibited strain-specific differences. Mice were administered either 0.05 mL mineral oil (n=5) or 0.05 mL 10% CCl_4_ diluted in mineral oil (n=5). After 3 days, animals were sacrificed and livers collected for future analysis as described.^4^

### Liver Tumors

HCC was chemically induced by injecting male B6C3F1 mice at p14 with either diethylnitrosamine (DEN, n=16) at a dose of 10 µL/g body weight or PBS (n=5) as control. The B6C3F1 mice were used since they develop robust HCC tumors after DEN treatment and spontaneous HCC tumors. Mice were maintained under normal conditions for 36 weeks at which time they were euthanized. Body and liver weights were determined, and visible tumors were counted, for each mouse. Liver tumors from DEN-injected mice and livers from age-matched PBS control mice were collected and stored at - 80°C until further use. Spontaneous HCC samples and age-matched control livers from male and female B6C3F1 mice were kindly provided by Dr. M. J. Hoenerhoff.^19^

### RNA extraction, cDNA synthesis, and Quantitative Real-Time PCR

Samples stored at -80° C were thawed on ice. Approximately 100 mg of tissue or 10^6^ cells were homogenized in 1 mL RNAzol RT (Sigma #R4533) and mRNA was extracted according to the product technical bulletin. cDNA was synthesized from 1 ug of RNA using the iScript cDNA Synthesis Kit (BioRad #170-8891). Reverse Transcriptase Quantitative PCR (RT-qPCR) reactions using SYBR Green PCR Supermix (BioRad #172-5275) were performed with a CFX96 Touch Real-Time PCR Detection System and results were analyzed with the CFX Manager software (BioRad). Oligonucleotide primer sequences are listed in Table 1. RT-qPCR C_T_ values for mouse studies were normalized to the indicated reference gene to evaluate relative tissue expression values. Reference genes were validated for stable expression in both control and experimental groups in each experiment. The 2^-ΔCT^ method was used to calculate expression fold changes.^20^

**Table 1.**
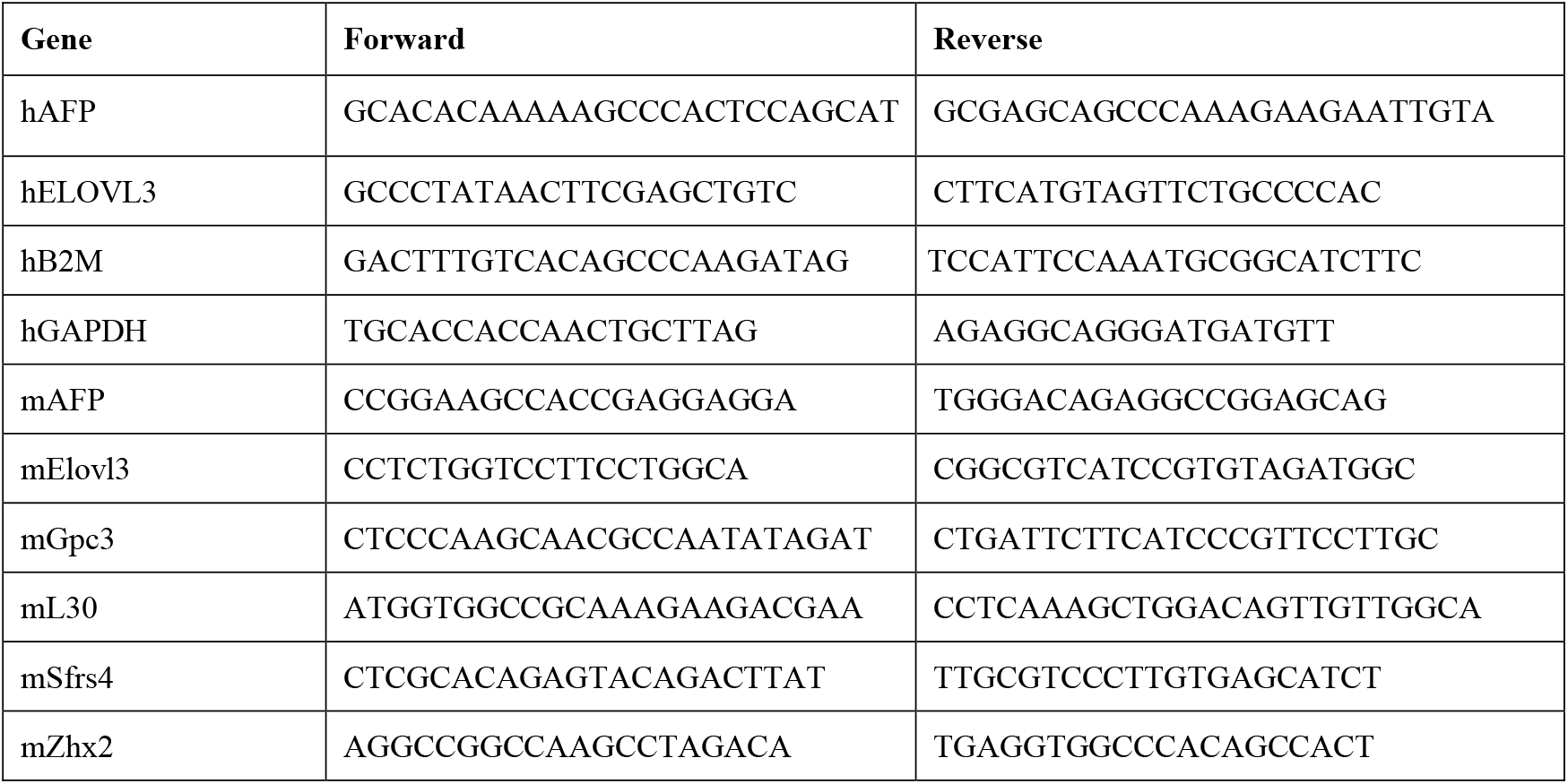
Primer sequences for RT-qPCR

### Western Blots

Western blots were prepared as described.^6,7^ Briefly, protein lysates (30µg/sample) were applied to SDS-PAGE gels, transferred to PVDF membranes, and probed with primary antibodies against the FLAG epitope (Sigma Cat. # F1804-1MG) and reprobed with a rabbit polyclonal anti-actin (Sigma Cat. # A2066) and species-specified HRP-conjugated secondary antibodies (Santa Cruz goat anti-mouse IgG-HRP (Cat # sc-2031) and anti-rabbit IgG-HRP (Cat # sc-2004)), followed by detection with the SuperSignal West Pico-Plus ECL kit (Thermo Scientific Cat. # 34580).

### Analysis of hepatic VLCFA

Hepatic VLCFA were analyzed by chemical derivatization and ultra-high-performance liquid chromatography tandem mass spectrometry (LC-MS/MS) using adaptations of the methods of Bollinger, *et al*.^21^ and carried out at the University of Kentucky Small Molecular Mass Spectrometry Core Laboratory. Liver samples (∼10 mg) were homogenized in 0.5 mL 0.1 M HCl, then 10 µl homogenate was mixed with 50 µl of internal standard (250 ng/mL Octadecanoic acid-d35), 0.5 mL 0.1 M HCl, 2 mL methanol and 1 mL chloroform. After vortex-mixing for 5 min, an additional 1 mL chloroform and 1.3 mL 0.1 M HCl were added, followed by an additional 5 min of vortexing, and the two phases were separated by centrifugation at 3000x*g* for 10 min. The lower phase was removed and evaporated to dryness under filtered nitrogen. Dried lipids were incubated with 1 mL 1 N KOH in ethanol at 90°C for 2 hours. After saponification, the solution was transferred to a 12-mL glass tube, the vial was rinsed with 1 mL ethanol, and the combined ethanol solution was extracted with 2 mL chloroform and 1.8 mL PBS. After phase separation, the lower layer was transferred to a 4 mL vial and dried under nitrogen. The dried material was derivatized with N-(4-aminomethylphenyl) pyridinium as described,^21^ followed by LC-MS/MS on the same day using an AB Sciex 4000 Q Trap coupled with an Exion LC system. The Analyst software package was used for data collection and analysis. Chromatography was carried out with a C_8_ reverse-phase column (Waters ACQUITY UPLC BEH C8, 2.1 × 50 mm, 1.7 µm) maintained at 40°C with the flow rate set to 0.4 mL/min. Solvent A was 100% water/ 0.1 % formic acid, and solvent B was acetonitrile/ 0.1% formic acid. A gradient program was used as follow (T min/%A): 0/90, 0.5/90, 0.51/80, 10.0/30, 10.1/0, 12.0/0, 12.1/90, 15.0 /90. The injection volume was 2.0 µl. The mass spectrometer was equipped with an electrospray ionization (ESI) source and operated in positive mode under the following operating parameters: IonSpray Voltage 5.5 kV, Desolvation temperature 50°C Ion Source Gas 1 40 psi, Ion Source Gas 2 50 psi, Curtain Gas 30 psi, Collision Gas Medium, Declustering Potential 160 V, Entrance Potential 10.0 V, and Collision Energy 55.0 V. Quantitative analysis was conducted by monitoring the precursor ion to production ion transition of each analyte.

### Expression Plasmids

A full-length expression vector for the 271 amino acid mouse ELOVL3 protein was generated by PCR amplification of mouse liver cDNA (Forward primer: GCCACCATGGACACATCCATGAATTTCTCAC; Reverse primer: GGATCCTTGGCTCTTGGATGCAACTTTG). Amplicons were cloned into the pGEM-T Easy vector (Promega #A1360), sequenced, excised using EcoRI and BamHI restriction enzymes, and cloned into the pcDNA3.1 expression vector (Invitrogen V790-20). *Elovl3* and empty vector pcDNA3.1 expression plasmids were transformed into competent E. coli cells using a standard cell transformation protocol. Plasmid preparations were performed using a Plasmid Max Kit (Qiagen #12165). The *Zhx2* expression plasmid was described previously.^7^

### Cell Culture and Transfection of Expression Plasmids

HepG2, Huh7, and HeLa cells were cultured in Dulbecco’s minimal eagle’s media (DMEM, Corning Cellgro #10-017-CV) supplemented with 10% fetal bovine serum (FBS, Fisher #03-600-511) and maintained at 37°C and 5% CO_2_. Cells were seeded onto 10 cm^2^ plates and cultured for 24 hours to obtain 70-80% confluence. Transfections using Turbofect (Thermo Scientific #R0531) were performed according to the manufacturer’s protocol. In some experiments, cells were co-transfected with a GFP expression plasmid to visualize transfection efficiency prior to further experimentation or for cell sorting as described below.

### Chromatin immunoprecipitation (ChIP)

ChIP analysis was performed as previously described.^7^ Briefly, adult male mouse livers from *Zhx2*^*WT*^ and *Zhx2*^*KO*^ mice were harvested and processed for ChIP with rabbit anti-Zhx2 antibodies (Bethyl labs) or control IgG using the Magna ChIP HiSense kit (Millipore). Primers were used to generate amplicons from the Elovl3 promoter and DNase hypersensitive sites (DHSs) located roughly 7 kb (3’ DHS) and 21 kb (5’ DHS) upstream of the Elovl3 promoter (oligonucleotide primers shown in Table 2). ChIP DNA samples were analyzed by quantitative PCR using Universal SYBR Green Supermix (Bio-Rad) and a Bio-Rad CFX96 real-time PCR system.

**Table 2.**
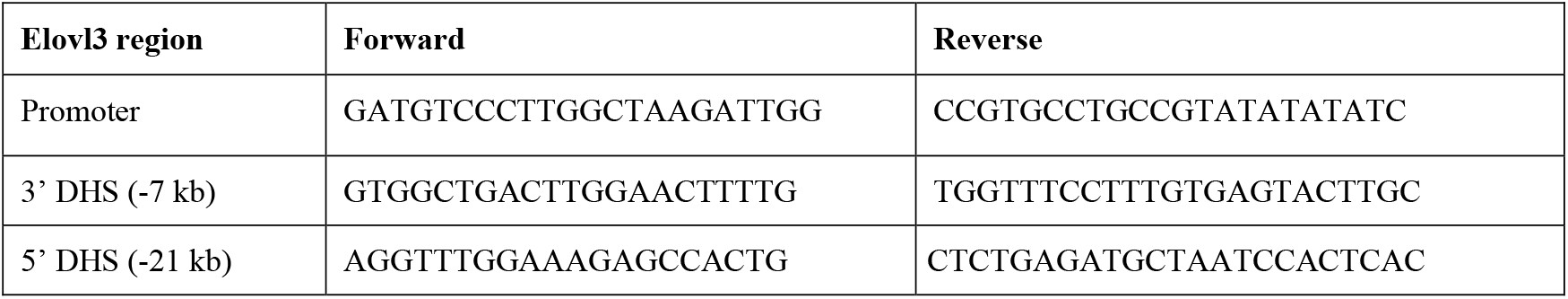
Primer sequences for ChIP

### Growth in Monolayers

Huh7 cells were seeded in 6-well plates and cultured for 24 hours and then transfected with either *Elovl3* or pcDNA expression plasmids. After 72 hours, cells were trypsinized, collected, and counted by hemocytometer with Trypan Blue dead cell exclusion.

### Growth in Soft Agar

Transfected Huh7 cells were cultured for 24 hours, then treated with trypsin to collect cells. Washed and resuspended cells (5 × 10^3^) were suspended in growth media and 0.8% agar, seeded between two layers of 1.2% agar mixed with growth media and then cultured for one week in 96-well plates. Cell viability was measured by CellGlo Titer Luminescent Cell Viability Assay (Promega #G7570) according to the product protocol and results were read on a luminometer. Sample luminescent values were normalized to wells containing media and agar layers without cells to account for background. Luminescent values for control cells were set to 100% and compared to values of *Elovl3* transfected cells in three separate experiments, with each sample assayed in technical triplicates.

### Cell Synchronization and Analysis of Cell Cycle

HeLa cells co-transfected with GFP and either *Elovl3* or *pcDNA3*.*1* expression plasmids were cultured for 24 hours then synchronized by blocking cell growth using a double Thymidine block and serum starvation.^22^ After release into fresh growth media, cells were immediately collected (T_0_) and at subsequent timepoints as designated. For cell cycle analysis, cells fixed overnight in 70% ethanol were stained with propidium iodide (Roche #11348639001) and filtered to remove cell aggregates immediately prior to analysis by Flow cytometry. Cells were gated for GFP expression to focus on transfected cells and then analyzed for DNA content to determine the percentage of cells in each cycle phase. FACS was performed by the University of Kentucky Flow Cytometry & Cell Sorting core facility.

### Statistical analysis

Data are graphed as mean +/- standard deviation. Statistical significance was determined by student’s t-test or ANOVA with a p-value <0.05 considered significantly different. Data were graphed and analyzed using GraphPad Prism software.

## Results

### Liver Elovl3 mRNA levels are reduced in the absence of Zhx2

The BALB/cJ mouse substrain carries a natural hypomorphic mutation in *Zhx2*, providing a model to identify putative ZHX2 target genes. Microarray data indicated that adult male liver *Elovl3* mRNA levels are reduced in BALB/cJ mice, suggesting that ZHX2 is required for normal *Elovl3* expression.^8^ To test the influence of ZHX2 on hepatic *Elovl3* expression, we analyzed *Elovl3* mRNA levels by RT-qPCR in adult male livers from BALB/cJ mice and the highly related BALB/cByJ substrain, which has wild-type *Zhx2* expression. *Elovl3* mRNA levels are substantially lower in BALB/cJ than in BALB/cByJ mice (Fig. 1A). In addition, since *Elovl3* is expressed in hepatocytes of the adult liver, male *Zhx2*^Δ*hep*^ mice, in which *Zhx2* is deleted solely in hepatocytes, and control littermates (*Zhx2*^*f/f*^) were examined for differences in *Elovl3* mRNA levels. In the absence of hepatocyte *Zhx2* expression, *Elovl3* mRNA levels are significantly reduced in male *Zhx2*^Δ*hep*^ mice (Fig. 1C), whereas *AFP*, which is repressed by ZHX2, is expressed at higher levels in *Zhx2*^Δ*hep*^ mice (Fig. 1D). These results are consistent with *Elovl3* regulation by ZHX2.

**Figure 1.**
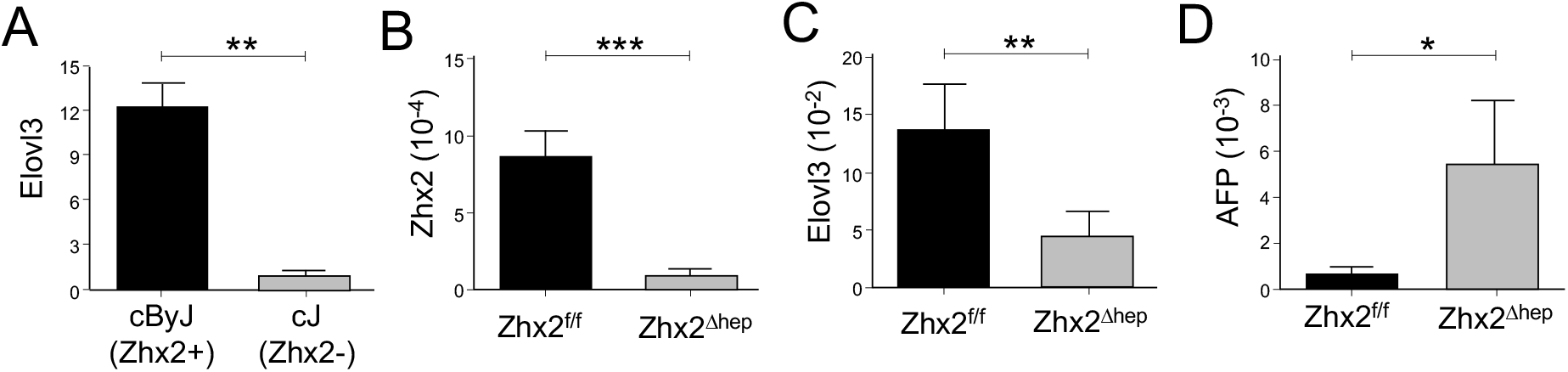
*Elovl3* mRNA levels are reduced in the absence of *Zhx2*. RNA was prepared from the livers of adult male **(A)** BALB/cByJ (cByJ; Zhx2+) and BALB/cJ (cJ; Zhx2-) mice and **(B-D)** BL/6 mice that express (*Zhx2*^*f/f*^) or do not express *Zhx2* in hepatocytes (*Zhx2*^Δ*hep*^). *Elovl3* (A,C), *Zhx2* (B) and *AFP* (D) levels were measured by RT-qPCR and normalized to *Sfrs4* (A) or ribosomal protein *L30* (B,C,D) mRNA levels. Mean (SD), analyzed by Student’s t-test, (n=3-5/group). *, p<0.05; **, p<0.01; *** p<0.001.

### Zhx2 activates Elovl3 expression in transfected HepG2 cells and binds putative Elovl3 control regions in adult liver

We next investigated whether ZHX2 controls *Elovl3* expression. Transient transfections were performed in human hepatoma HepG2 cells, which normally exhibit very low *ELOVL3* mRNA levels. Cells were transiently transfected with a *Zhx2* expression vector or empty vector control. After 48 hours, cells were harvested and mRNA and protein were prepared. Western blot analysis confirmed increased Flag-ZHX2 protein levels in transfected cells (Fig. 2A). The *Elovl3* mRNA levels were markedly higher in *Zhx2*-transfected cells compared to control cells, supporting our mouse data indicating that ZHX2 is a positive regulator of *Elovl3* expression. In contrast, *AFP*, which is expressed at high levels in HepG2 cells and is repressed by ZHX2, exhibited reduced expression in *Zhx2*-transfected cells (Fig. 2B).

**Figure 2.**
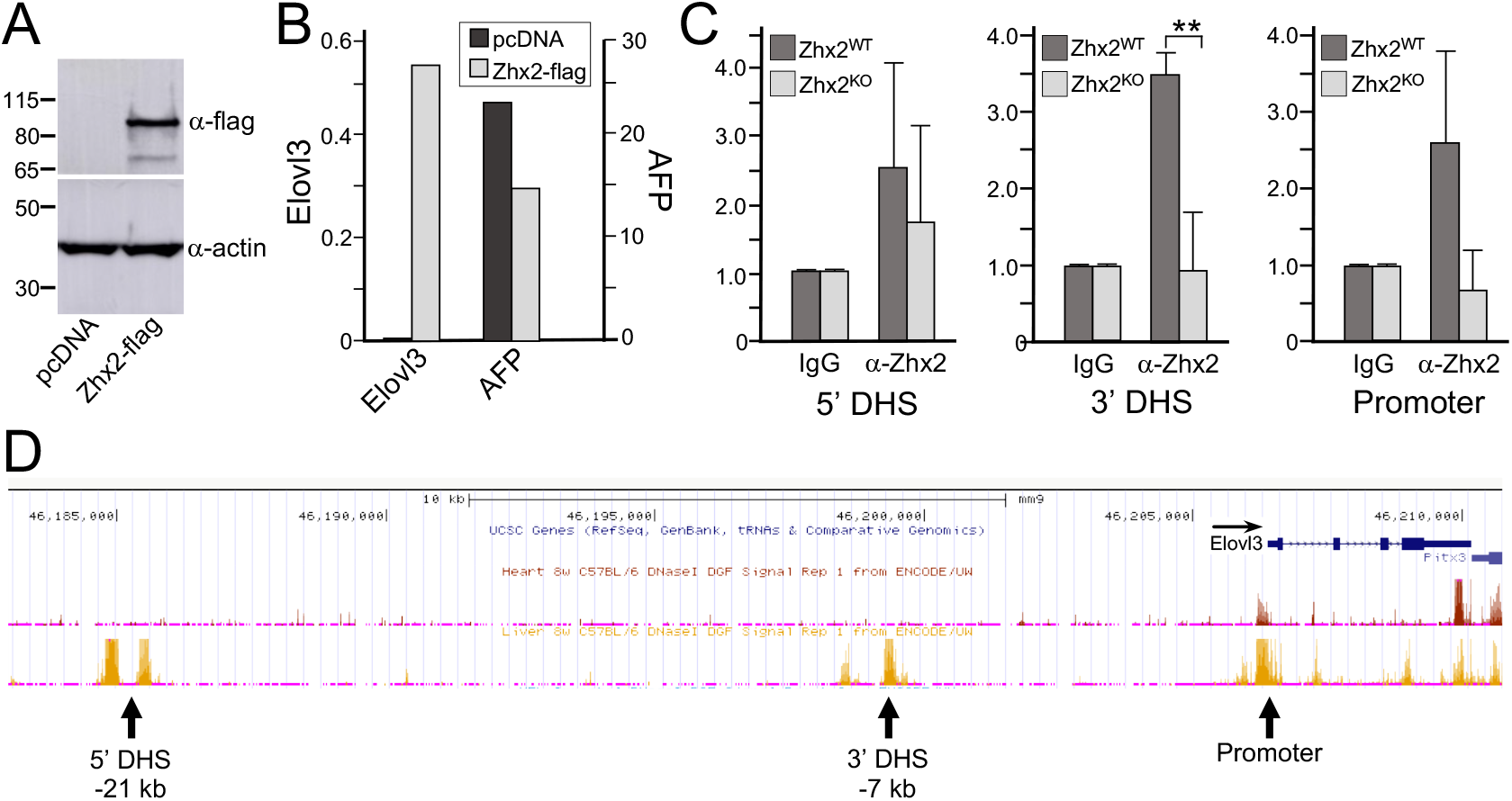
*Zhx2* activates Elovl3 expression in transfected HepG2 cells and binds putative Elovl3 control regions in adult liver. **(A, B)**. Human hepatoma HepG2 cells were transiently transfected with a pcDNA vector containing flag-tagged Zhx2 (Zhx2-flag) or pcDNA empty vector control (pcDNA). After 48 hours, cells were harvested and RNA and protein extracts were prepared. **(A)** Western blots were probed using anti-Flag antibodies and reprobed with anti-actin antibodies to confirm equal lysate loading. **(B)** RT-qPCR analysis was performed to measure *Elovl3* and *AFP* mRNA levels normalized to *GAPDH*. Data is representative of three separate experiments. **(C)**. Adult male liver samples from *Zhx*^*WT*^ and *Zhx2*^*KO*^ mice were analyzed by ChIP using anti-Zhx2 antibodies or control IgG and primers that generate amplicons from the Elovl3 promoter, 3’ DHS site or 5’ DHS site. For all three regions, levels of IgG in *Zhx2*^*WT*^ and *Zhx2*^*KO*^ mice were set to 1. Data are from three separate experiments. **, p<0.01 (D). DNAse Hypersensitive sites from ENCODE adult mouse liver flanking the mouse Elovl3 gene are shown as yellow peaks (designated by arrows) in analysis using the University of California – Santa Cruz (UCSC) Genome Browser.^41^

To test whether ZHX2 directly binds to regions that control *Elovl3* expression, ChIP experiments were performed with adult liver samples from *Zhx2*^*WT*^ or *Zhx2*^*KO*^ mice with anti-ZHX2 antibodies or control IgG. Although studies have identified several genes that are directly regulated by ZHX2, including AFP, Cyclin D1, and Mup20, a consensus ZHX2 binding site has not been identified. Furthermore, *Elovl3* regulatory elements have not been characterized. Therefore, to identify candidate ZHX2 binding regions controlling *Elovl3* expression, we searched for DHSs surrounding the *Elovl3* gene using the ENCODE database. This analysis identified DHSs at the *Elovl3* promoter and regions that are ∼7 kb (3’ DHS) and ∼21 kb (5’ DHS) upstream of the *Elovl3* exon 1 (Fig. 2D). Using primers to amplify these three regions, ChIP analysis from *Zhx2*^*WT*^ mice indicated that ZHX2 was enriched in the 3’ DHS and *Elovl3* promoter (although binding to the promoter did not reach statistical significance; *p* = 0.07), but not in the 5’ DHS site (Fig. 2C). No enrichment of these regions were found with anti-ZHX2 antibodies when liver samples from *Zhx2*^*KO*^ mice were used. These data suggest that ZHX2 directly activates *Elovl3* expression.

### Male-biased Elovl3 expression in the liver is established postnatally

A number of previously described ZHX2 targets, including those that are sex-biased, exhibit changes in expression in the perinatal liver.^1,6, 7^ We characterized developmental changes in liver *Elovl3* expression in male and female mice. In male mice, *Zhx2* mRNA levels increase gradually between embryonic day 17.5 (e17.5) and postnatal day 28 (p28) and then have a nearly 7-fold increase between p28 and p56 (Fig. 3A). A very similar pattern is seen with *Elovl3*, with levels barely detectable at e17.5, showing a modest gain between p14 and p21, followed by a greater than 6-fold increase between p28 and p56 (Fig. 3B). A different pattern of *Zhx2* and *Elovl3* occurs in female liver. *Zhx2* mRNA levels increase slightly between p7 and p14, followed by a greater increase between p21 and p28 (Fig. 3C). At p56, *Zhx2* levels in male and female livers are nearly the same. Female hepatic *Elovl3* mRNA levels are low but detectable at e17.5 and remain relatively constant until p21, followed by a slight increase at p28, then a steep decline between p28 and p56 (Fig. 3D). In contrast, *AFP* expression exhibits a consistent, dramatic decline in the postnatal liver in both male and female mice.^7^ Thus, hepatic *Elovl3* expression is fully activated in males and nearly completely repressed in females between 4 and 8 weeks of age, establishing the male-biased expression pattern that persists in the healthy liver of adult mice.

**Figure 3.**
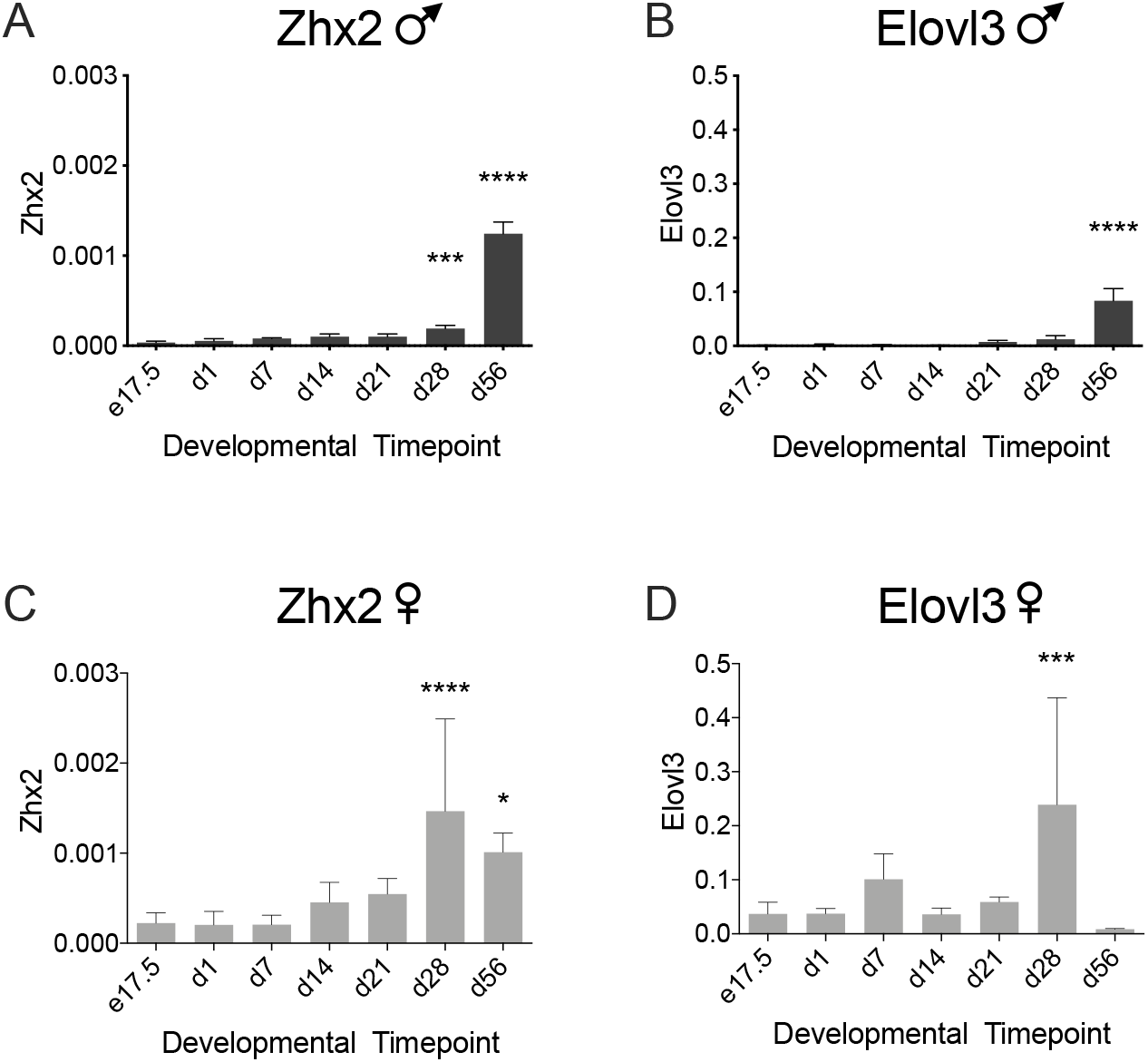
Male-biased *Elovl3* expression in the adult liver is established between four and eight weeks of age. Liver mRNA was prepared from male and female B6C3F1 mice at embryonic day 17.5 (e17.5) and postnatal day 1 (p1), p7, p14, p21, p28 and p56. RT-qPCR analysis was performed to assess *Zhx2* (panels A and C) and *Elovl3* (panels B and D) mRNA levels in male (panels A and B) and female (panels C and D) mice, which were normalized to *L30*. Mean (SD), analyzed by ANOVA with Dunnett’s test for multiple comparisons; each timepoint compared to e17.5 expression. n= 4-10 mice/group. *, <0.05; ***, <0.001; ****, <0.0001.

### Elovl3 mRNA levels and ELOVL3-synthesized lipids decrease in regenerating liver

Expression of many ZHX2 target genes become dysregulated in liver disease. In general, genes that are repressed by ZHX2 in the postnatal liver, including *AFP, Gpc3*, and *H19*, are activated during liver regeneration and in liver tumors.^1,4,23,24^ Conversely, genes that are postnatally activated by ZHX2, including certain *Mups* and *Cytochrome p450 4a12* (*Cyp4a12*), exhibit decreased levels in regenerating livers and tumors.^6,7^ We therefore assessed *Elovl3* expression in multiple models of liver damage and disease. Liver regeneration in response to acute injury is a process in which normally quiescent hepatocytes reenter the cell cycle to replace damaged cells that are lost due to injury and restores normal liver mass and function.^25^ To monitor changes in *Elovl3* expression during liver regeneration, the CCl_4_ model was used.^26^ Adult male C3H and BL/6 mice were administered a single i.p. injection of either CCl_4_ or mineral oil (M.O.) as control and sacrificed after 72 hours, when regeneration is robust. In both mouse strains, hepatic *Elovl3* mRNA levels were reduced in CCl_4_-treated compared to control mice, although the decrease is more significant in C3H mice (Fig. 4A). In contrast to *Elovl3, AFP mRNA* levels increased in both strains three days after CCl_4_ treatment; this increase is greater in C3H than in BL/6 mice (Fig. 4B). We then assessed whether decreased *Elovl3* expression during regeneration was reflected in hepatic lipid composition. In the liver, VLCFAs are synthesized by various elongase enzymes with some redundancy, but also with a great deal of specificity for carbon chain length and saturation (Fig. 4C). We analyzed free fatty acids and saponified lipids extracted from B6C3F1 mouse livers, which have robust *Elovl3* repression and *AFP* induction similarly to C3H mice (data not shown), and found that regenerating livers have significantly lower concentrations of C22:1 monounsaturated and C20:0-C22:0 saturated VLCFAs (Fig. 4D-E, G-H). There were no differences in C20:1, C24:0, or C24:1 VLCFAs (Fig. 4F, I). Further, there was a notable difference in liver lipid concentration depending on carbon length and desaturation: multiple lipid saturated and monounsaturated species C22 and shorter were decreased during regeneration, whereas C22 and longer polyunsaturated VLCFAs were generally increased (Table 3).

**Table 3.** Saponified Lipids in MO and CCl4-treated livers

| (ug/mg liver) | MO | CCl4 | Effect in CCl4 | Significance |
| --- | --- | --- | --- | --- |
| C14:0 | 0.7097 | 0.6340 | Down |  |
| C16:0 | 5.6696 | 4.8819 | Down |  |
| C18:0 | 5.3628 | 4.0986 | Down | * |
| C18:1 | 4.2443 | 3.6730 | Down |  |
| C18:2 | 5.7281 | 5.4713 | Down |  |
| C19:0 | 0.3068 | 0.2354 | Down | * |
| C20:0 | 0.3565 | 0.0978 | Down | * |
| C20:1 | 1.6373 | 1.1540 | Down |  |
| C20:2 | 1.6404 | 1.7800 | Up |  |
| C20:3 | 2.0457 | 1.9632 | Down |  |
| C20:4 | 5.4518 | 5.9013 | Up |  |
| C20:5 | 0.6718 | 0.7168 | Up |  |
| C21:0 | 0.0103 | 0.0061 | Down | * |
| C22:0 | 0.0827 | 0.0453 | Down | * |
| C22:1 | 0.2925 | 0.1498 | Down | * |
| C22:2 | 0.1133 | 0.1159 | No Change |  |
| C22:3 | 0.2253 | 0.2889 | Up |  |
| C22:4 | 1.2871 | 2.3613 | Up | * |
| C22:5 | 1.0816 | 1.5071 | Up | * |
| C22:6 | 2.2403 | 2.7369 | Up | * |
| C23:0 | 0.0116 | 0.0112 | No Change |  |
| C24:0 | 0.0200 | 0.0182 | Down |  |
| C24:1 | 0.2870 | 0.3540 | Up |  |
| C24:2 | 0.0647 | 0.1594 | Up | * |
| C24:3 | 0.0146 | 0.0455 | Up | * |
| C24:4 | 0.2159 | 0.5545 | Up | * |
| C24:5 | 0.1565 | 0.2751 | Up | * |
| C24:6 | 0.1358 | 0.1806 | Up |  |
| C26:1 | 0.0017 | 0.0036 | Up | * |
| C26:2 | 0.0023 | 0.0070 | Up | * |
| C26:3 | 0.0061 | 0.0058 | No Change |  |
| C26:4 | 0.0075 | 0.0279 | Up | * |
| C26:5 | 0.0114 | 0.0437 | Up | * |
| C26:6 | 0.0147 | 0.0274 | Up | * |
| C34:0 | 0.0035 | 0.0019 | Down |  |
| C36:2 | 0.0025 | 0.0015 | Down |  |

**Figure 4.**
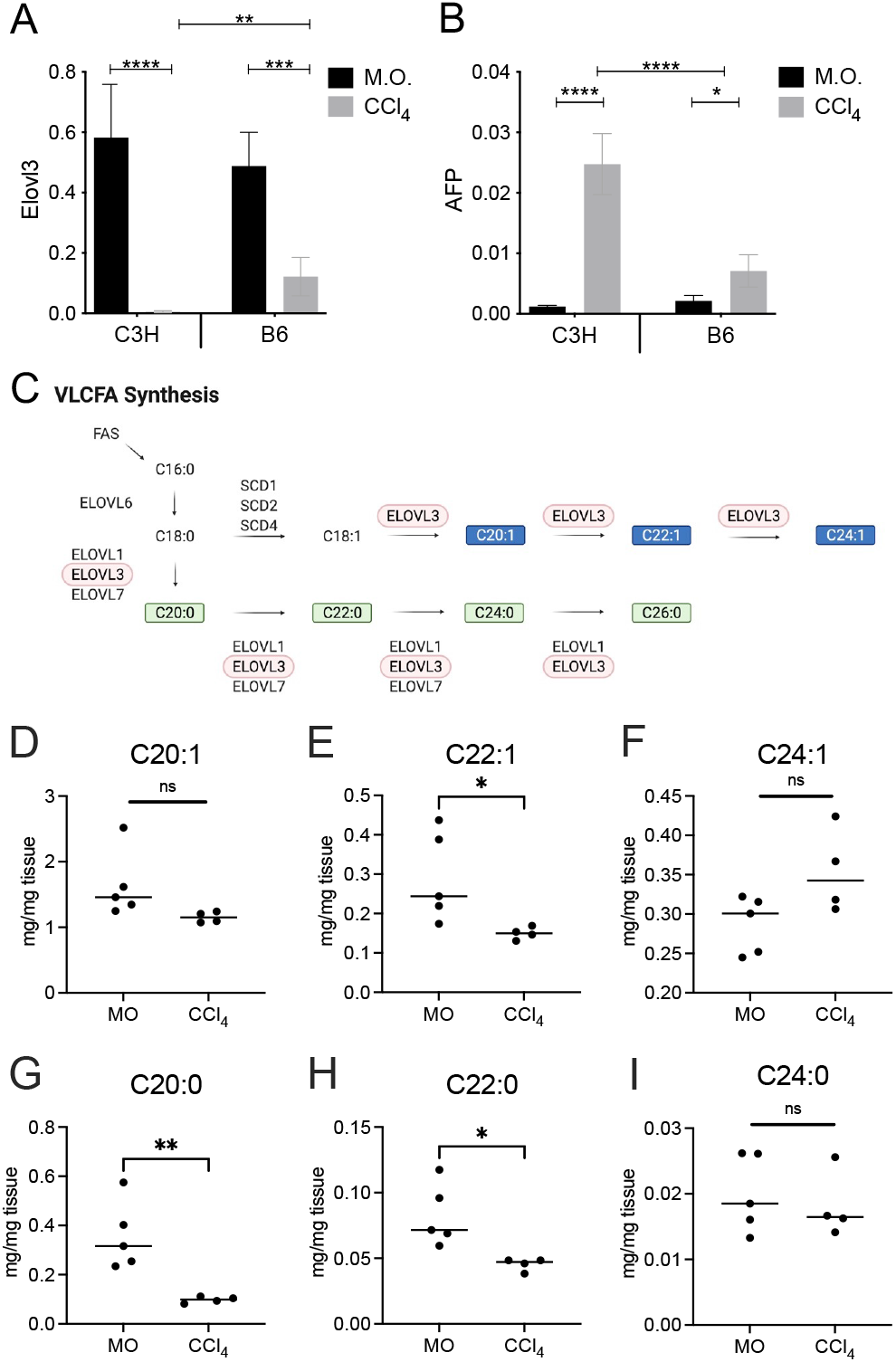
*Elovl3* mRNA levels and ELOVL3-synthesized lipids decrease in regenerating liver. **(A, B)** Adult male C3H or BL/6 mice were injected with vehicle (mineral oil, M.O.; black bars) or CCl_4_ (gray bars) and sacrificed after 3 days to assess mRNA changes during liver regeneration (n=5/group). RT-qPCR was performed to analyzed hepatic *Elovl3* (panel A) and *AFP* (panel B) mRNA levels in both strains. **(C)** Schematic of VLCFA *de novo* synthesis pathway highlighting ELOVL3 metabolites, modified from Guillou, et al.^40^ Those in blue and green represent monounsaturated and saturated VLCFA, respectively. **(D-I)** Liver lipids were extracted and analyzed for lipid composition by LC-MS/MS and normalized to the weight of the tissue mass. Data is shown for VLCFAs synthesized by ELOVL3; C20:1 (D), C22:1 (E), C24:1 (F), C20:0 (G), C22:0 (H) and C24:0 (I). Data for all analyzed VLCFAs are shown in Table 3. Mean (SD), A, B analyzed by 2-way ANOVA with Tukey correction for multiple comparisons, D-I analyzed by Student’s t-test; * p<0.05; ** p<0.01, *** p<0.001, ****, p<0.0001.

### Elovl3 mRNA is decreased in liver tumors and forced Elovl3 expression diminishes tumor cell viability slows cell growth

The dramatic decrease in *Elovl3* mRNA levels in regenerating livers led us to test whether *Elovl3* levels also change in HCC in two mouse models. First, the well-established DEN mouse model of HCC was used.^27^ DEN was administered to p14 mice to initiate and propagate chemically-induced HCC tumors, which were harvested after 36 weeks. The liver/body weight ratios in DEN treated mice and number of visible tumors were significantly greater in DEN-treated mice than in PBS-treated age-matched controls, consistent with previous DEN tumor models (Fig. 5A, B).^27^ In addition, *Gpc3*, a known HCC marker in mice and humans, is higher in tumors of DEN mice than in PBS-treated controls (Fig. 5C). Significantly, *Elovl3* mRNA levels were dramatically lower in tumor samples compared to age-matched PBS-injected control littermates (Fig. 5D). In a second tumor model, we examined liver mRNA from control livers and spontaneous HCC tumors in aged B6C3F1 male and female mice.^19^ The male-biased *Elovl3* expression was again apparent in the livers of control mice (Fig. 5B). In both sexes, *Elovl3* levels were nearly undetectable in HCC tumors, with a much more pronounced reduction observed in male mice since Elovl3 mRNA levels were already very low in female control livers (Fig. 5B).

**Figure 5.**
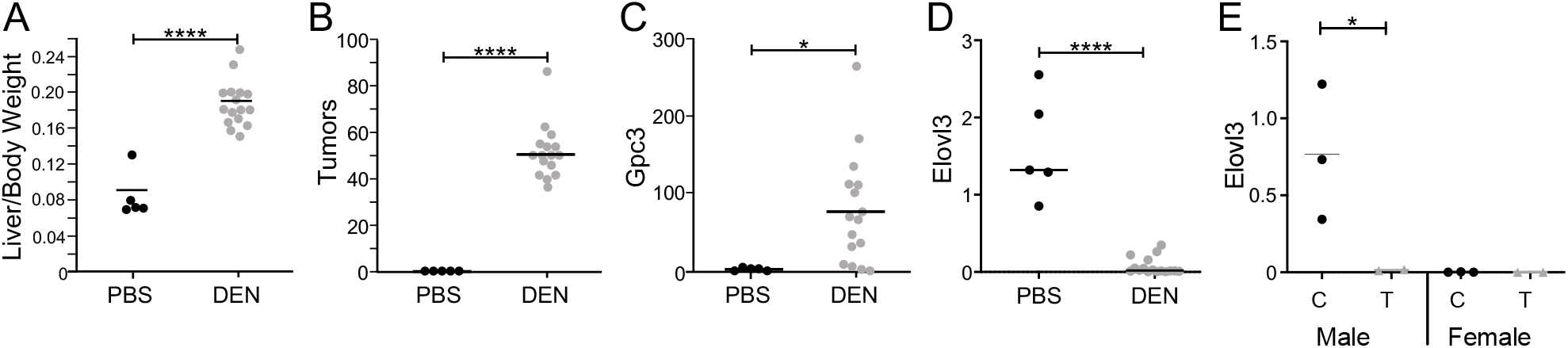
*Elovl3* mRNA levels are decreased in liver tumors. **(A-D)** HCC was initiated in male B6C3F1 mice by administering DEN (n=16) at p14; control mice (n=5) with were injected with PBS. Livers were harvested 36 weeks post-injection. Liver/body weight ratios **(A)** and the number of visible tumors **(B)** were determined for each mouse. RNA was prepared from control livers or tumors from each mouse and *Gpc3* **(C)** and *Elovl3* **(D)** mRNA levels were determined. **(E)** Spontaneous HCC tumors (T) were removed from B6C3F1 mice (tumors from 3 males, 2 females) that were approximately 2 years old; normal control livers (C) were removed from age-matched controls (3 male, 2 female). In both sets of cohorts, *Elovl3* or *Gpc3* mRNA levels were measured by RT-qPCR and normalized to *Sfrs4*. Mean (SD), analyzed by Student’s t-test; * p<0.05; ****, p<0.0001.

### Increased Elovl3 expression reduces cell proliferation and inhibits the S-G2 transition

Previous studies demonstrated that ZHX2 inhibits the growth of liver tumor cell lines, implicating a role for ZHX2 target genes in the association between ZHX2 and HCC.^10,28,29^ Since *Elovl3* expression is nearly absent in HCC tumors, we evaluated whether increased *Elovl3* expression alone would influence tumor cell growth. Human HCC Huh7 cells, which have negligible endogenous *Elovl3* expression, were plated as monolayers and transfected with an *Elovl3* expression vector or an empty vector control (pcDNA), and viable cells were counted after 72 hours. In four independent experiments, *Elovl3* overexpression reduced the number of cells by roughly 45% (Fig. 6A). Anchorage-independent growth is a signature feature of oncogenesis that can be gauged by cellular growth and expansion in soft agar. Huh7 cells transfected with either the *Elovl3* expression vector or pcDNA empty vector were seeded in media-supplemented soft agar, cultured for one week, and measured using a luminescent viability assay. In three independent experiments, *Elovl3* transfected cells exhibited a 34% reduction in cell viability compared to pcDNA3.1-transfected control cells (Figure 6B). Taken together, these data indicate that elevated *Elovl3* levels diminish Huh7 cell viability.

**Figure 6.**
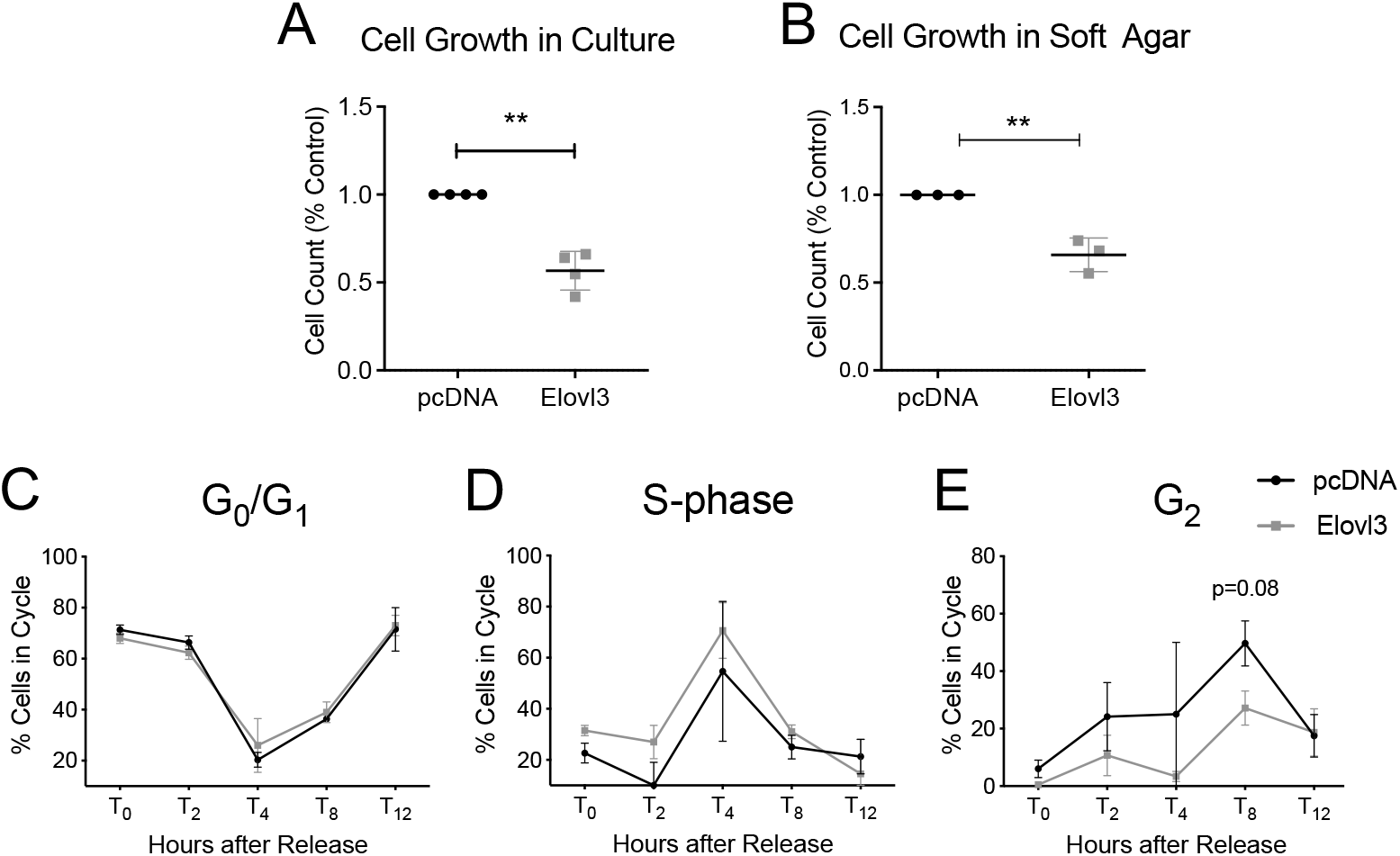
*ELOVL3* overexpression reduces Huh7 cell growth *in vitro* and stalls Hela cells at the S-G2 transition. **(A-B)**. Cells were transfected with a pcDNA empty vector control (pcDNA) or an *Elovl3* expression vector (*Elovl3*). **(A)** Huh7 cells were maintained in culture as monolayers for 72 hours then washed, trypsinized, and viable cells were counted. (**B**) Twenty-four hours after transfection, Huh7 cells were trypsinized, seeded in media-supplemented agar and cultured for 1 week. Cell viability was measured using a luminescence assay. In both sets of experiments, pcDNA-transfected cell counts were set to 1. Monolayer cultures were performed in 4 independent experiments; soft agar growth repeated in 3 independent experiments. Mean (SD), analyzed by Student’s t-test; ** p<0.01. **(C-E)**. Hela cells co-transfected with GFP and either human *ELOVL3* expression plasmid or empty vector were synchronized in G_0_ by double Thymidine block in serum-free media, then released into growth media. Cells gated for GFP expression were analyzed by FACS for cell cycle stage (G_0_/G_1_; S-phase; G_2_) by Propidium iodide staining at the designated timepoints.

To further explore the effect of *Elovl3* expression on cell cycle progression, highly proliferative HeLa cells were co-transfected with a GFP expression plasmid along with either the *Elovl3* expression vector or control pcDNA and then synchronized in G_0_. After release into growth media, cells were collected immediately or at multiple timepoints for up to 12 hours and stained with propidium iodide. Using flow cytometry, GFP-positive cells were analyzed for DNA content. As early as 2 hours after release there was a notable increase in the proportion of *Elovl3*-transfected cells in S-phase and a concomitant decrease in the G_2_ cell population compared to control transfected cells (Fig. 6C-E). This trend continued to hold for the first 8 hours after release.

## Discussion

ZHX2 is required for the complete postnatal repression of numerous fetal liver genes, including *AFP, H19* and *Gpc3*; the continued expression of these genes in the adult liver of BALB/cJ mice is due to a natural mutation in the *Zhx2* gene in this strain.^1,4^ In contrast to this repression, ZHX2 is required for the in the developmental activation of *Mup* genes and certain *Cyp* genes, both of which also exhibit sex-specific regulation.^6,7^ Here, we demonstrate that ZHX2 is a positive regulator of *Elovl3* in the adult male mouse liver. *Elovl3* mRNA levels are lower in BALB/cJ mice compared to the BALB/cByJ substrain, which has a wild-type *Zhx2* gene, and in adult BL/6 mice with a hepatocyte-specific *Zhx2* deletion compared to wild-type littermate controls (Fig. 1A-D). Consistent with these data, *Zhx2* transfections in HepG2 cells significantly increase endogenous *Elovl3* mRNA levels (Fig. 2). We also show that *Elovl3* mRNA levels are significantly reduced in regenerating liver and in both DEN-induced and spontaneous HCC (Figs 4, 5); a concordant decrease in lipids synthesized by ELOVL3 were found in regenerating livers (Fig. 4, D-I). Our data indicate that elevated *Elovl3* expression diminishes the growth of Huh7 cells and stalls HeLa cell in S-phase (Fig. 6), suggesting that decreased *Elovl3* expression in HCC may enable increased cellular growth and proliferation.

*Elovl3*, like other ZHX2 targets, is developmentally regulated, and as with other known sex-specific ZHX2 target genes, male-biased *Elovl3* expression becomes apparent between 4-8 weeks of age (Fig. 3). In fact, *Elovl3* is expressed higher in female liver during earlier developmental stages (Fig. 3B, D). The mechanism by which *Elovl3* decreases in female and increases in male livers is not known, although these changes occur concurrently with sexual maturation in mice that occurs 4-8 weeks after birth. Previous studies in castrated mice demonstrated that androgens are required for male-biased hepatic *Elovl3* expression, and that female mice injected with 5α-dihydrotestosterone exhibited increased *Elovl3* expression with circadian oscillations that mimicked male expression patterns.^14,30^ It is not clear whether cross-talk between ZHX2 and androgens is required for *Elovl3* regulation, but our data support that ZHX2 directly binds the *Elovl3* promoter and an upstream DHS to regulate transcription. Additional studies are needed to clarify the relationship between ZHX2, steroid hormone metabolism, and male-biased *Elovl3* expression.

*Zhx2* target genes *AFP, Gpc3*, and *H19* are silenced at birth but transiently reactivated in regenerating liver after a single CCl_4_ treatment.^4,23,24,31^ The extent of this induction is less in BL/6 mice than in other strains, including C3H.^24^ This strain-specific difference is a monogenic trait that has been mapped to a locus on mouse Chromosome 2 called *alpha-fetoprotein regulator 2 (Afr2*).^31^ In an opposite expression pattern compared to these other ZHX2 targets, *Elovl3* levels increase postnatally in male mice (Fig. 3B) and are reduced regenerating liver (Fig. 4A). Interestingly, this *Elovl3* silencing is less robust in BL/6 mice than in C3H liver (Fig. 4A), suggesting *Afr2* also impacts *Elovl3* repression in regenerating liver. The product of the *Afr2* locus has not been identified, but the fact that ZHX2 targets also appear to be controlled by *Afr2* during liver regeneration suggests an interaction between ZHX2 and *Afr2* in the adult liver.

Hepatocyte proliferation is robust during embryonic development, slows in the perinatal period and remains low in the healthy adult liver; liver regeneration reestablishes transient and rapid hepatocyte proliferation to replace damaged and dead cells.^32^ Conversely, *Elovl3* levels are extremely low in embryonic livers, increase postnatally (Fig. 3B), and decrease dramatically during liver regeneration (Fig. 4A). This expression pattern supports the possibility that *Elovl3* may inhibit cellular proliferation and that its downregulation permits cellular expansion. Consistent with this, we find that *Elovl3* overexpression slows the *in vitro* growth of Huh7 cells (Fig. 5A, B) and stalls HeLa cell cycle progression in S-phase (Fig. 6B-C). Our data from DEN-induced and spontaneous HCC models suggest that reduced *Elovl3* levels may also influence proliferation of transformed cells (Fig. 5A, B). The notion that *Elovl3* influences cell cycle regulation is also supported by a study demonstrating that *Elovl3* is transcriptionally activated by the tumor suppressor p53, which has a well-established role in cell cycle arrest.^33^ Our data suggest that VLCFAs synthesized by ELOVL3 contribute to its regulation of cell proliferation. The C20 to C24 saturated and monounsaturated VLCFA synthesized by ELOVL3 are often incorporated into ceramides and other classes of bioactive lipids. Ceramides are effective tumor suppressors by inducing apoptosis and reducing proliferation through cell cycle arrest.^34-36^ Reduced *Elovl3* expression (Fig. 4A), resulting in lower hepatic C20-C22 VLCFAs levels, are seen in regenerating liver (Fig. 4D-I) may reduce certain bioactive lipids and allow increased cellular proliferation. More detailed metabolic tracing will be required to fully elucidate the lipid species and direct effects of ELOVL3-synthesized VLCFAs on proliferation and tumorigenesis and to investigate the potential role of other elongases in these processes. ELOVL7 was found to contribute to prostate cancer by increasing production of saturated VLCFA that were synthesized into androgen steroids, serving as a potent prostate cancer cell growth signal.^37^ Interestingly, *Elovl7* mRNA was also increased in our models of HCC (B.T.S., unpublished observation). Future studies characterizing and tracing the lipid species produced by ELOVL3 and other elongases will provide insight into this family of enzymes and their effects on cell proliferation.

Both tissue culture and mouse xenograft studies indicate that ZHX2 functions as a tumor suppressor in the liver, presumably through its control of target genes.^10,11,38,39^ The ZHX2 target genes *AFP, H19 and Gpc3* serve as valuable HCC biomarkers but their roles in tumorigenesis are not fully understood. ZHX2 directly represses the transcription of Cyclins A and E to inhibit cell cycle progression.^10^ Recently, ZHX2 was shown to repress hepatic *Lpl* and SREBP1c levels in human liver cell lines, which inhibited lipid uptake and de novo synthesis, respectively, and stalled xenograft tumor growth.^11^ Data provided here identify *Elovl3* as another link between ZHX2 and tumor suppression, potentially through the synthesis of C20-C:22 VLCFAs. Additional studies will be needed to further elucidate the role of ZHX2 and its target genes, including ELOVL3, in liver physiology and disease, including HCC.

## Abbreviations

AFP: alpha-fetoprotein
Afr2: alpha-fetoprotein regulator 2
CCl_4_: Carbon tetrachloride
Cyp: Cytochrome P450
DEN: diethylnitrosamine
DHS: DNase hypersensitive site
Elovl3: Elongation of Very Long Chain Fatty Acids 3
Gpc3: Glypican 3
HCC: hepatocellular carcinoma
Lpl: lipoprotein lipase
M.O.: mineral oil
Mup: major urinary protein
RT-qPCR: Reverse Transcriptase-quantitative PCR
VLCFA: very long chain fatty acids
ZHX2: Zinc fingers and homeoboxes 2

## Data Availability Statement

The data that support the findings of this study are provided in the results section and described in the Methods and Materials. Additional details and primary data are available on request from the corresponding author.

## Author Contributions

K.T. Creasy contributed to experimental design and conduct, data interpretation and manuscript writing, H. Ren contributed to experimental design and conduct and data interpretation, J. Jiang contributed to experimental conduct and data interpretation, M.L. Peterson and B.T. Spear contributed to experimental design, data interpretation and manuscript writing.

## Acknowledgments

We thank Shirley Qiu for helpful discussions and technical assistance, Andrew Morris and Jianzhong Chen of the University of Kentucky Small Molecular Mass Spectrometry Core Laboratory for VLCFA analysis and Mark Hoenerhoff (University of Michigan) for spontaneous liver tumor samples.

## References

1. Perincheri S, Dingle RW, Peterson ML, Spear BT. Hereditary persistence of alpha-fetoprotein and H19 expression in liver of BALB/cJ mice is due to a retrovirus insertion in the Zhx2 gene. Proc Natl Acad Sci, USA 2005; 102:396–401.

2. Perincheri S, Peyton DK, Glenn M, Peterson ML, Spear BT. Characterization of the ETnII-alpha endogenous retroviral element in the BALB/cJ Zhx2 (Afr1) allele. Mamm Genome 2008; 19:26–31.

3. Clinkenbeard EL, Turpin C, Jiang J, Peterson ML, Spear BT. Liver size and lipid content differences between BALB/c and BALB/cJ mice on a high-fat diet are due, in part, to Zhx2. Mamm Genome 2019; 30:226–236.

4. Morford LA, Davis C, Jin L, Dobierzewska A, Peterson ML, Spear BT. The oncofetal gene glypican 3 is regulated in the postnatal liver by zinc fingers and homeoboxes 2 and in the regenerating liver by alpha-fetoprotein regulator 2. Hepatology 2007; 46:1541–1547.

5. Peterson ML, Ma C, Spear BT. Zhx2 and Zbtb20: Novel regulators of postnatal alpha-fetoprotein repression and their potential role in gene reactivation during liver cancer. Semin Cancer Biol 2011; 21:21–27.

6. Creasy KT, Jiang J, Ren H, Peterson ML, Spear BT. Zinc Fingers and Homeoboxes 2 (Zhx2) Regulates Sexually Dimorphic Cyp Gene Expression in the Adult Mouse Liver. Gene Expression 2016; 17:7–17.

7. Jiang J, Creasy KT, Purnell J, Peterson ML, Spear BT. Zhx2 (zinc fingers and homeoboxes 2) regulates major urinary protein gene expression in the mouse liver. J Biol Chem 2017; 292:6765–6774.

8. Gargalovic PS, Erbilgin A, Kohannim O, Pagnon J, Wang X, Castellani L, Leboeuf R, Peterson ML, Spear BT, Lusis AJ. Quantitative trait locus mapping and identification of Zhx2 as a novel regulator of plasma lipid metabolism. Circ Cardiovasc Genet 2010; 3:60–67.

9. Lv Z, Zhang M, Bi J, Xu F, Hu S, Wen J. Promoter hypermethylation of a novel gene, ZHX2, in hepatocellular carcinoma. Amer J Clin Path 2006; 125:740–746.

10. Yue X, Zhang Z, Liang X, Gao L, Zhang X, Zhao D, Liu X, Ma H, Guo M, Spear BT, Gong Y, Ma C. Zinc fingers and homeoboxes 2 inhibits hepatocellular carcinoma cell proliferation and represses expression of Cyclins A and E. Gastroenterology 2012; 142:1559–1570.

11. Wu Z, Ma H, Wang L, Song X, Zhang J, Liu W, Ge Y, Sun Y, Yu X, Wang Z, Wang J, Zhang Y, Li C, Li N, Gao L, Liang X, Yue X, Ma C. Tumor suppressor ZHX2 inhibits NAFLD-HCC progression via blocking LPL-mediated lipid uptake. Cell Death Differ 2020; 27:1693–1708.

12. Jakobsson A, Westerberg R, Jacobsson A. Fatty acid elongases in mammals: their regulation and roles in metabolism. Progress Lipid Res 2006; 45:237–249.

13. Hannun YA, Obeid LM. Principles of bioactive lipid signalling: lessons from sphingolipids. Nat Rev Mol Cell Biol 2008: 9:139–150.

14. Brolinson A, Fourcade S, Jakobsson A, Pujol A, Jacobsson A. Steroid hormones control circadian Elovl3 expression in mouse liver. Endocrinology 2008; 149:3158–3166.

15. Tvrdik P, Asadi A, Kozak LP, Nedergaard J, Cannon B, Jacobsson A. Cig30, a mouse member of a novel membrane protein gene family, is involved in the recruitment of brown adipose tissue. J Biol Chem 1997; 272:31738–31746.

16. Westerberg R, Tvrdik P, Unden AB, Mansson JE, Norlen L, Jakobsson A, Holleran WH, Elias PM, Asadi A, Flodby P, Toftgard R, Capecchi MR, Jacobsson A. Role for ELOVL3 and fatty acid chain length in development of hair and skin function. J Biol Chem 2004; 279:5621–5629.

17. Michelott, GA, Machado MV, Diehl AM. NAFLD, NASH and liver cancer. Nat Rev Gastroenterol Hepatol 2013; 10:656–665.

18. Tessari P, Coracina A, Cosma A, Tiengo A. Hepatic lipid metabolism and non-alcoholic fatty liver disease. Nutr Metab Cardiovasc Dis 2009; 19:291–302.

19. Hoenerhoff MJ, Pandiri AR, Lahousse SA, Hong HH, Ton TV, Masinde T, Auerbach SS, Gerrish K, Bushel PR, Shockley KR, Peddad, SD, Sills RC. Global gene profiling of spontaneous hepatocellular carcinoma in B6C3F1 mice: similarities in the molecular landscape with human liver cancer. Toxicologic Pathology 2011; 39:678–699.

20. Livak KJ, Schmittgen TD. Analysis of relative gene expression data using real-time quantitative PCR and the 2(-Delta Delta C(T)) Method. Methods 2001; 25:402–408.

21. Bollinger JG, Naika GS, Sadilek M, Gelb MH. LC/ESI-MS/MS detection of FAs by charge reversal derivatization with more than four orders of magnitude improvement in sensitivity. J Lipid Res 2013; 54:3523–3530.

22. Tong J, Sun D, Yang C, Wang Y, Sun S, Li Q, Bao J, Liu Y. Serum starvation and thymidine double blocking achieved efficient cell cycle synchronization and altered the expression of p27, p53, bcl-2 in canine breast cancer cells. Res Vet Sci 2016; 105:10–14.

23. Pachnis V, Belayew A, Tilghman SM. Locus unlinked to α-fetoprotein under the control of the murine raf and Rif genes. Proc Natl Acad Sci USA 1984; 81:5523–5527.

24. Belayew A, Tilghman SM. Genetic analysis of α-fetoprotein synthesis in mice. Mol Cell Biol 1982; 2:1427–1435.

25. Michalopoulos GK, DeFrances MC. Liver regeneration. Science 1997; 276:60–66.

26. Taniguchi M, Takeuchi T, Nakatsuka R, Watanabe T, Sato K. Molecular process in acute liver injury and regeneration induced by carbon tetrachloride. Life Sci 2004; 75:1539–1549.

27. Heindryckx F, Colle I, Van Vlierberghe H. Experimental mouse models for hepatocellular carcinoma research. Int J Experimental Pathology 2009; 90:367–386.

28. Luan F, Liu P, Ma H, Yue X, Liu J, Gao L, Liang X, Ma C. Reduced nucleic ZHX2 involves in oncogenic activation of glypican 3 in human hepatocellular carcinoma. Int J Biochem Cell Biol 2014; 55:129–135.

29. Shen H, Luan F, Liu H, Gao L, Liang X, Zhang L, Sun W, Ma C. ZHX2 is a repressor of alpha-fetoprotein expression in human hepatoma cell lines. J Cell Mol Med 2008; 12:2772–2780.

30. Chen H, Gao L, Yang D, Xiao Y, Zhang M, L, C, Wang A, Jin Y. Coordination between the circadian clock and androgen signaling is required to sustain rhythmic expression of Elovl3 in mouse liver. J Biol Chem 2019; 294:7046–7056.

31. Jin DK, Feuerman MH. Genetic mapping of Afr2 (Rif): regulator of gene expression in liver regeneration. Mamm Genome 1998; 9:256–258.

32. Spear BT, Jin L, Ramasamy S, Dobierzewska A. Transcriptional control in the mammalian liver: liver development, perinatal repression, and zonal gene regulation. Cell Mol Life Sci 2006; 63:2922–2938.

33. Kon N, Wang D, Li T, Jiang L, Qiang L, Gu W. Inhibition of Mdmx (Mdm4) in vivo induces anti-obesity effects. Oncotarget 2018; 9:7282–7297.

34. Dbaibo GS, Pushkareva MY, Jayadev S, Schwarz JK, Horowitz JM, Obeid LM, Hannun YA. Retinoblastoma gene product as a downstream target for a ceramide-dependent pathway of growth arrest. Proc Natl Acad Sci USA 1995; 92:1347–1351.

35. Lee JY, Bielawska AE, Obeid LM Regulation of cyclin-dependent kinase 2 activity by ceramide. Exp Cell Res 2000; 261:303–311.

36. Zhu XF, Liu ZC, Xie BF, Feng GK, Zeng YX. Ceramide induces cell cycle arrest and upregulates p27kip in nasopharyngeal carcinoma cells. Cancer Lett 2003; 193:149–154.

37. Tamura K, Makino A, Hullin-Matsuda F, Kobayashi T, Furihata M, Chung S, Ashida S, Miki T, Fujioka T, Shuin T, Nakamura Y, Nakagaw, H. Novel lipogenic enzyme ELOVL7 is involved in prostate cancer growth through saturated long-chain fatty acid metabolism. Cancer Res 2009; 69:8133–8140.

38. Kawata H, Yamada K, Shou Z, Mizutani T, Yazawa T, Yoshino M, Sekiguchi T, Kajitani T, Miyamoto K. Zinc-fingers and homeoboxes (ZHX) 2, a novel member of the ZHX family, functions as a transcriptional repressor. Biochem J 2003; 373:747–757.

39. Weng MZ, Zhuang PY, Hei ZY, Lin PY, Chen ZS, Liu YB, Quan ZW, Tang ZH. ZBTB20 is involved in liver regeneration after partial hepatectomy in mouse. Hepatob Pancreat Dis 2014; 13:48–54.

40. Guillou H, Zadravec D, Martin PG, Jacobsson A. The key roles of elongases and desaturases in mammalian fatty acid metabolism: Insights from transgenic mice. Progress Lipid Res 2010; 9:186–199.

41. Kent WJ, Sugnet CW, Furey TS, Roskin KM, Pringle TH, Zahler AM, Haussler D. The human genome browser at UCSC. Genome Research 2002; 12:996–1006.

